# Finding cell-specific expression patterns in the early Ciona embryo with single-cell RNA-seq

**DOI:** 10.1101/197699

**Authors:** Garth R. Ilsley, Ritsuko Suyama, Takeshi Noda, Nori Satoh, Nicholas M. Luscombe

## Abstract

Single-cell RNA-seq has been established as a reliable and accessible technique enabling new types of analyses, such as identifying cell types and studying spatial and temporal gene expression variation and change at single-cell resolution. Recently, single-cell RNA-seq has been applied to developing embryos, which offers great potential for finding and characterising genes controlling the course of development along with their expression patterns. In this study, we applied single-cell RNA-seq to the 16-cell stage of the *Ciona* embryo, a marine chordate and performed a computational search for cell-specific gene expression patterns. We recovered many known expression patterns from our single-cell RNA-seq data and despite extensive previous screens, we succeeded in finding new cell-specific patterns, which we validated by in situ and single-cell qPCR.

## Introduction

Cell types can be characterised at high resolution with single-cell RNA-seq. Apparently homogenous groups of cells can be clustered to identify novel and rare subtypes^1–5^ and cells undergoing differentiation at different rates can be ordered and analysed^6–9^. This offers great potential for studying developing embryos with their high diversity of cell types^10–20^. Making sense of this diversity is a goal of developmental biology and an important first step is identifying and characterising the relatively few genes controlling the course of development and the subsets of cells they are expressed in, or in other words, their gene expression patterns.

We applied single-cell RNA-seq (scRNA-seq) to the 16-cell stage of *Ciona*, a simple chordate, and looked for gene expression patterns using a novel computational approach. Zygotic expression begins around the 8-cell stage^21^ and many gene expression patterns, both maternal and zygotic, are known from comprehensive in situ screens in *Ciona*^22^ (Fig. 1b, Supplementary Table 1). We identified many of these from our scRNA-seq data. In addition, we found unknown gene expression patterns, which we validated by in situ and single-cell qPCR.

**Figure 1:**
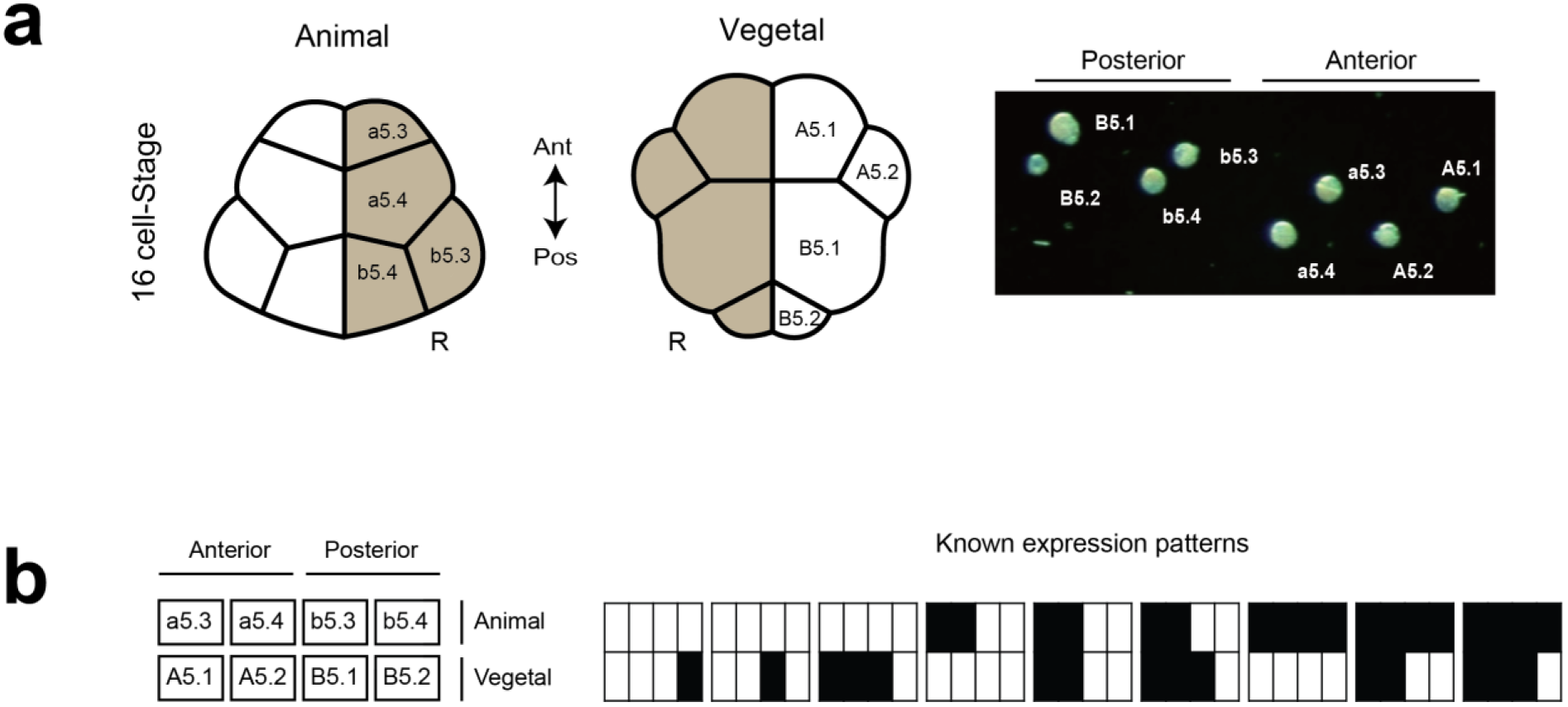
The cell types and gene expression patterns of the 16-cell *Ciona* embryo are well characterised. **(a)** Drawing of a 16 cell–stage embryo showing *Ciona* cell type nomenclature (left). All eight cells were dissected and isolated by glass needle from the right side of four 16-cell embryos, and identified visually under microscope (right). **(b)** Nine known expression patterns of *Ciona* (right) are shown according to a schematic layout (left), with solid rectangles indicating expression in the corresponding cells.

## Results

*Ciona* development follows an invariant lineage allowing cells to be unambiguously identified at the 16-cell stage^23–25^, whether before sequencing as in our work (Fig. 1a) or afterwards based on expression, as in recent work of the same stage^26^. We collected cells from four 16-cell stage embryos, each from different individuals fertilised on different days. The left and right side of the embryo at this stage are thought to be symmetrical, but to avoid any potential bilateral variation in our small sample, we collected all eight cell types from the right half of each embryo. We multiplexed the cells of each embryo (or batch) and sequenced them on Illumina MiSeq, as well as on three lanes across two Illumina HiSeq 2500 runs, leading to scRNA-seq measurements for 32 cells (Supplementary Table 2).

Our cells were sequenced with relatively high coverage (around 20 million reads per cell on Illumina HiSeq and 4.5 million per cell on MiSeq). Although we had only four replicates per cell type, we produced a reliable dataset.

### Assessment of technical variability and reproducibility

Our results show limited technical variation within each batch: the expression levels in different cell types from the same embryo are well correlated for embryos 2, 3 and 4. They are, in fact, more similar to each other than the same cell types are across different individuals (Fig. 2a-b). Although we cannot separate out the sources of cross-embryo variation, this result is consistent with a previous report showing that maternal mRNA levels vary significantly between unfertilized eggs from different individuals^27^. It is also worth noting that very little of the variation between embryos is from the sequencing run. This can be seen by comparing our sequence results from MiSeq with HiSeq—the correlation between unnormalised counts from the two platforms is over 99% for every cell type, whether genes with zero counts are included or excluded. This is consistent with previous results showing high correlation between expression measurements from tens of millions of reads per cell and those from lower coverage of a million or fewer reads^28,29^.

**Figure 2:**
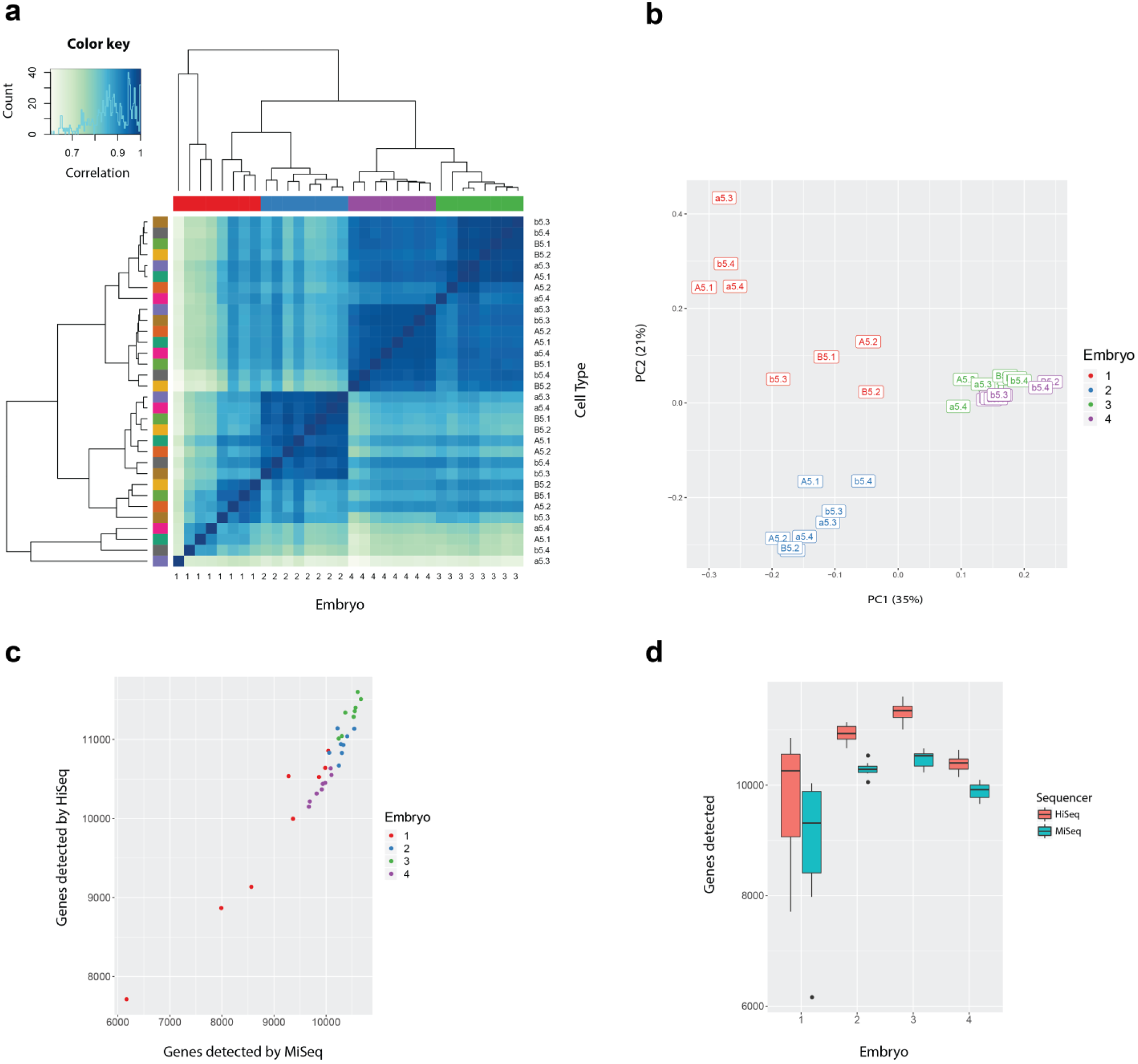
Our dataset exhibits low technical variability within batches and between Illumina MiSeq and HiSeq. **(a)** Clustered heat map of the symmetrical correlation matrix of normalised expression data from HiSeq samples (excluding ERCC counts and genes with zero counts), with the histogram in the top left providing the colour key for the correlation matrix. The embryo and cell type annotations for each sample are shown on different axes, but the samples are arranged in the same order. The samples are colour-coded by embryo along the top and by cell type on the left with the same hierarchical clusters shown in both cases. (**b**) PCA plot of the same data showing the first two components, which explain 56% of the total variance. The samples are colour-coded by embryo and labelled by cell type. (**c and d**) The number of genes detected is consistent across the four embryo replicates whether the libraries were sequenced on MiSeq or HiSeq. (**c**) Scatter plot showing the consistent relationship between MiSeq and HiSeq, with more zeros or undetected genes for some cells of embryo 1 compared to the others. (**d**) Boxplots showing the narrower distribution of genes detected for embryos 2 to 4 compared to embryo 1 and the consistent increase from MiSeq to HiSeq.

This embryo batch effect is further demonstrated by a Principal Components Analysis (Fig. 2b), which shows a similar result with the cell types of embryos 2, 3 and 4 being close to each on the first two components (which explain 56% of the variance) and the cell types of embryo 1 being more spread out.

The close clustering of cells from the same embryo, as well as their high correlation, suggests that our technical variability is low, leading to reproducibility within each batch (or embryo). A confirmation of the reproducibility of our results is the tight distribution of genes detected across samples within embryos (Fig. 2c-d). Genes were considered detected when the measured count was greater than zero. These results show that slightly more genes were detected on HiSeq than MiSeq, but that the median difference for each embryo is less than 10%. This can be compared with a previous result showing a reduction of genes detected of around 39% when lowering sequence coverage to less than a million reads per cell^28^. As before, embryo 1 showed more variability across samples than the other embryos.

Further, our data can be compared to previously published data for the 16-cell stage that was generated using gene expression microarrays from many pooled single cells^27^. We found good agreement with our scRNA-seq expression patterns for the key genes shown in their paper (Supplementary Fig. 1).

### Normalisation of counts

After producing a dataset of counts, we normalised our data. Since we were interested in gene expression differences between different cell types, we did not normalize across genes (such as by GC content or transcript length), but only for sequencing depth by dividing by the total number of reads per sample. More complex normalisation steps could be considered^30^, but this simple approach suffices for our data. It gives a natural measure of expression for each gene, namely the proportion it contributes to the total, which we assume is independent of the total number of reads.

We then made use of a suitable transformation of proportions, the arcsine of the square root (referred to here as the φ transformation), namely

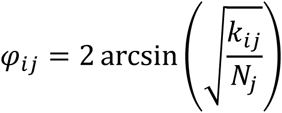

where *k_ij_* is the count for the i^th^ gene and j^th^ sample and N_j_ is the total number of counts for the j^th^ sample. The difference between φ_i_ of two samples can be interpreted as an effect size index for proportions, namely Cohen’s h^31^.

In practice, the arcsine function in the φ transformation is largely redundant because most genes are expressed at low proportions. At these low values, the arcsine transformation is close to identity, meaning that a square root transformation of proportions performs equivalently in many cases.

### Finding putative gene expression patterns

We then took a simple and effective approach to find gene expression patterns. Instead of grouping cells into a single set of clusters, we clustered the gene expression pattern of each gene separately. For each gene, we grouped the different cell types into two classes of ON and OFF expression. We took advantage of our known cell types and calculated the Euclidean distance between vectors of replicate measurements for each cell and performed single-linkage hierarchical clustering of these. The resulting top-level clusters determined the ON/OFF pattern of each cell type for each gene.

We then ranked these results, by taking an approach that does not require parametric estimation of variation or dispersion. We calculated our cluster reliability score as the difference between the first quartile of the ON cluster and the third quartile of the OFF cluster, which we call the Transquartile Range (TQR). The TQR is larger when the difference in cluster means is larger, but it penalizes higher variation for a given difference in means.

### Ranking cell-specific gene expression patterns

We applied this approach and produced a list of ranked genes and their expression patterns. We examined our results for 77 genes (Supplementary Table 1) with known in situ patterns (Imai et al. 2004; Matsuoka et al. 2013; Miwata et al. 2006; Prodon et al. 2007). Our top 35 results matched the known in situ patterns (Supplementary Fig. 2a), except for KH.L152.12, which is validated below as a new pattern by in situ and qPCR (Fig. 4). However, the lower ranked results did not correspond well (Supplementary Fig. 2b).

There were a few reasons for this. For example, clustering did not produce a correct pattern for *Lefty* (Supplementary Fig. 2b). The cell types, A5.2 and B5.1, are normally considered part of the *Lefty* pattern, but they have intermediate expression in our scRNA-seq results and, in this instance, the clustering algorithm places them in the OFF cluster. For a few other genes, e.g. *DPOZ* (KH.C12.589) and *Dlx.b*, the clustering is correct, but the TQR is low. For a few other genes, no reads were mapped, e.g. *Sox7/17/18*, *Fringe 2* and KH.C13.22, but in most cases where our scRNA-seq data does not agree with published in situ patterns, our expression measurements were low or relatively uniform across the eight cells and thus the algorithm functions correctly in attributing lower score to these results. Many of these genes are expressed maternally in the *Ciona* embryo and hence are not easily detectable by our RNA-seq protocol, which amplifies from the poly(A) tail of mRNA. In situ hybridisation can detect localized maternal signal in the cytoplasm, which could explain the discrepancy. It is also possible that some in situ results are false positives.

### Validating cell-specific gene expression patterns

Using these known in situ results, we assessed that the top 40 ranked results from all genes were likely to be reliable and focused on these for further validation. We found 12 distinct patterns in the top 40, which included all known patterns as well as three potentially new patterns (highlighted in orange in Fig. 3). Ten patterns are currently known at this stage in *Ciona*^27, 32–37^, and although we matched only nine of these directly (Fig. 1b), the tenth pattern has a single known case, *Tfap2-r.b* (*AP-2-like2*), which we did pick up, but without expression in A5.2. This agrees with previous observations that expression is not consistent in this cell across embryos^27,35^. Further, it is in agreement with the average over many embryos as measured by microarray^27^ (Supplementary Fig. 1).

**Figure 3:**
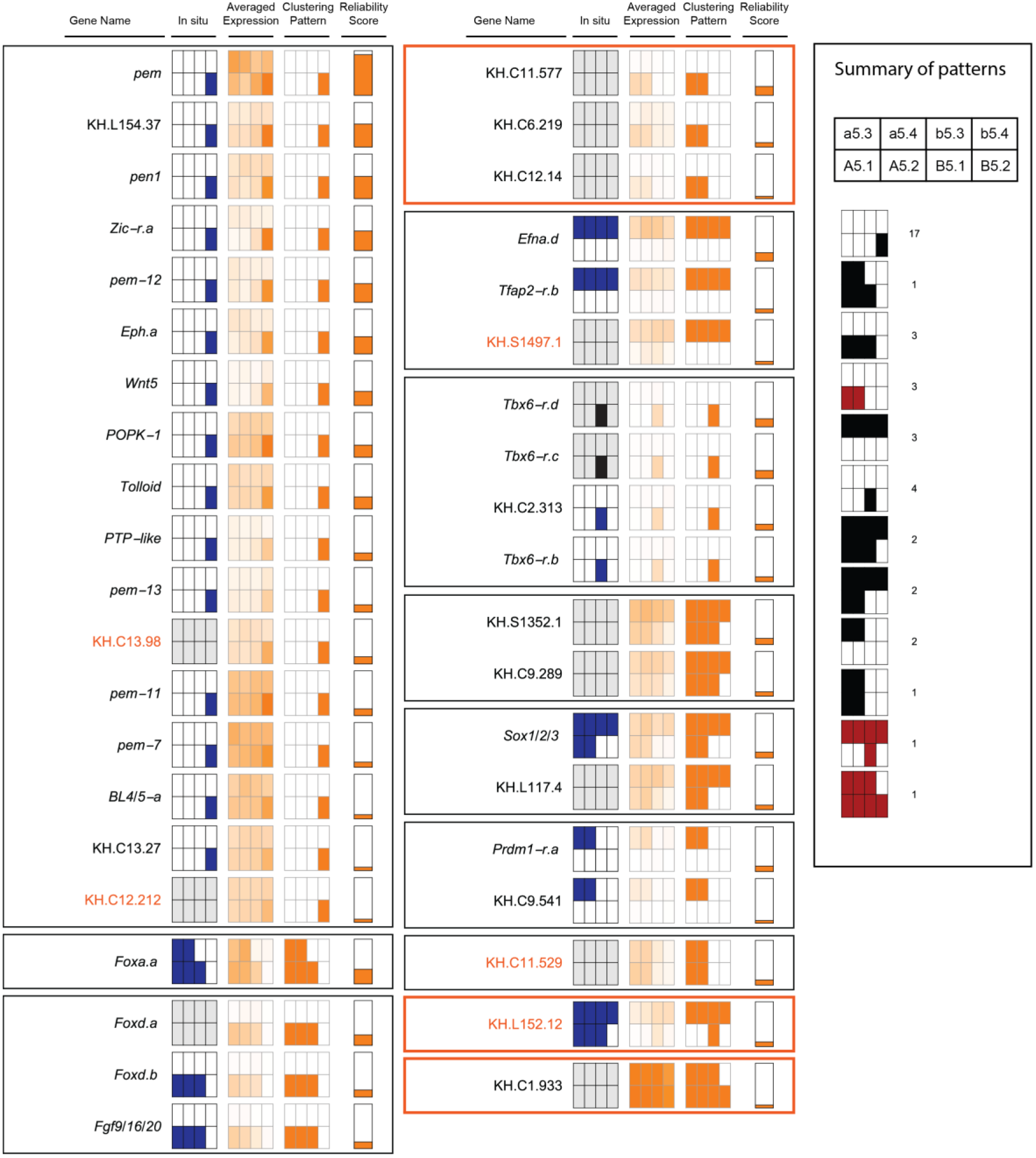
All previously known patterns occur in the top 40 genes when ranked according to their Transquartile Range (TQR). Expression summary plots of the top 40 genes, grouped by pattern, with an orange border indicating an unknown pattern. The arrangement of cells in each pattern is shown in the summary box on the right, together with the number of genes per pattern, whether known (black) or unknown (red). Genes whose patterns we validated are highlighted in orange. For each gene, the columns indicate the gene name, any previously known in situ pattern in blue (grey if not known), the average of the transformed expression values (clipped above 0.05), the pattern resulting from clustering, and finally, the TQR as the reliability score, scaled for visualization. The in situ probe for Tbx6-r.b also detects Tbx6-r.c and Tbx6-r.d (indicated by a black in situ pattern).

We validated one of the potentially new patterns by in situ and single-cell qPCR, namely the pattern for KH.L152.12 (Fig. 4e). We could not validate the second pattern for KH.C1.933 that has relatively ubiquitous expression, although with apparently reduced expression in b5.4. We also did not manage to validate the third potentially new pattern (expression in A5.1 and A5.2 only), although the scRNA-seq expression data in this case seems to show clear expression in these cells.

**Figure 4:**
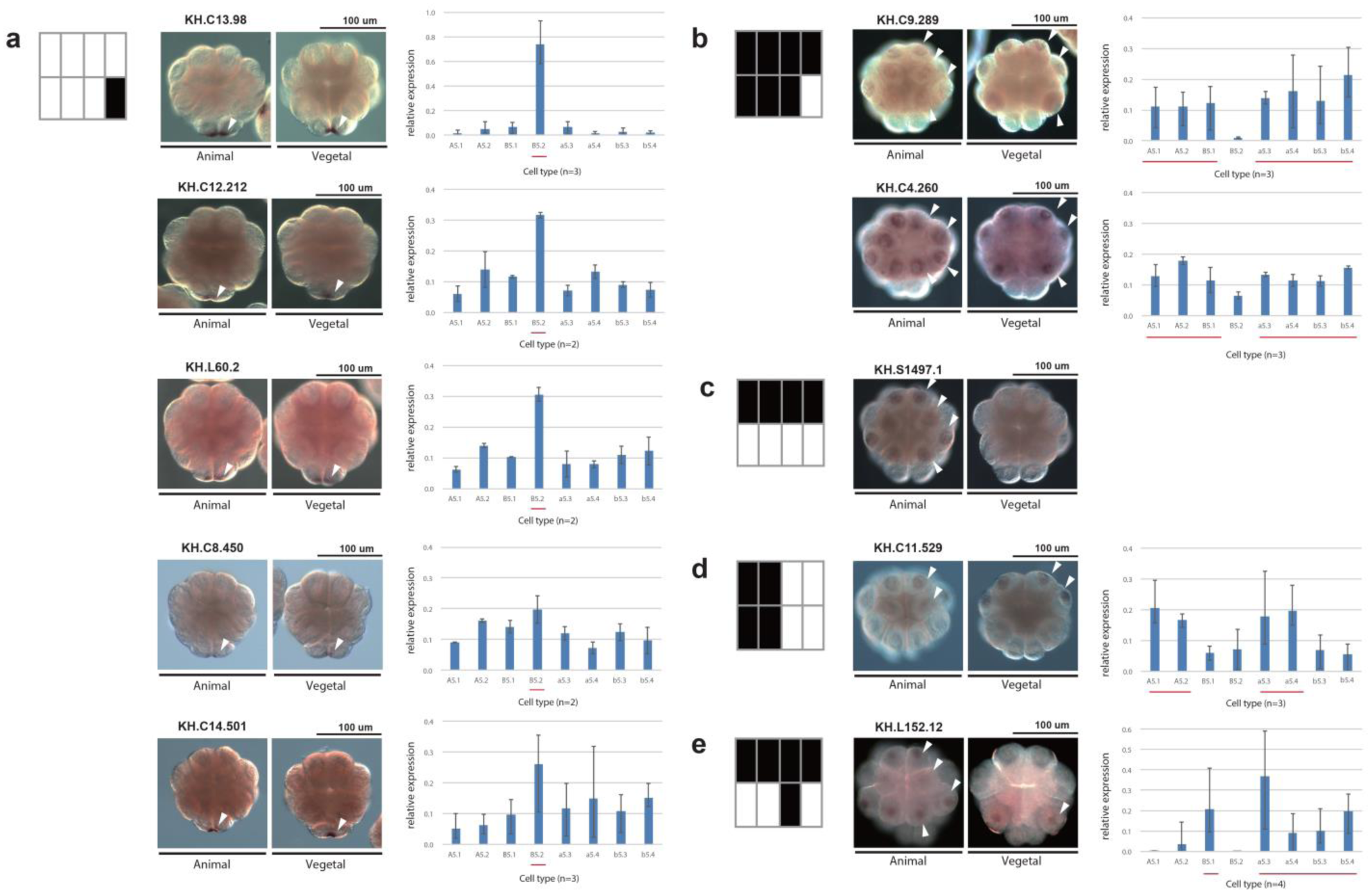
Pattern discovery results are validated by in situ hybridisation and single-cell qPCR. (**a-e**) For each pattern being tested (left), the in situ hybridisation results are shown from the animal and vegetal side (middle). The arrowheads pointing to the expressing cells are shown on only one side of the embryo. The scale bar indicates 100 μm. Gene expression levels for each cell type were measured by single-cell qPCR. The qPCR measures for the cell types of each embryo were scaled to add to 1. The bar charts (right) show the mean of the resulting values of biological replicates, with whiskers indicating the range. The lines under the bars indicate the cells of the expression pattern being tested. The number of replicates of each gene is shown under each bar chart.

In addition to recovering all known patterns and validating one novel pattern, we found at least 28 genes with known in situ patterns. Of the 12 patterns we found, the pattern with expression in the B5.2 cell only was the most represented. This is also the most frequent pattern in known in situ patterns, i.e. postplasmic/PEM RNAs^38^. The majority of our results for B5.2 are confirmed by previous in situ datasets, but we identified new B5.2-specific genes, such as KH.C13.98 and KH.C12.212, confirming their expression by in situ and single-cell qPCR (Fig. 4a).

We also validated other classes of uncharacterized genes, namely KH.S1497.1, which expresses specifically in the animal hemisphere, and KH.C11.529 on the anterior side (Fig. 4c-d). These results are particularly striking, since it was expected that no further zygotic genes expression patterns would be found at this stage in *Ciona*^22^.

We also looked more widely in the top 60 (Supplementary Fig. 3) and validated additional genes, KH.C8.450, KH.C9.289 and KH.C4.260, by single-cell qPCR and in situ hybridisation (Fig. 4b). The first is another example of B5.2 expression, whereas the last two genes are expressed in all cells except B5.2, a pattern known previously from *Hes.a*^39^. While developing our approach, we also applied the TQR to the microarray data of this stage^27^ and looked for reliable B5.2 cell-specific expression using both datasets. As a result, we also found and validated KH.L60.2 and KH.C14.501 (ranked 88 and 2403 in our data; see Supplementary Table 7).

In conclusion, we have recovered many known patterns, as well as patterns and genes that had not been detected previously despite extensive in situ screens. These results open up opportunities for further research into developmental patterning in *Ciona*. In addition, we have demonstrated that single-cell RNA-seq is a viable alternative to extensive in situ screens, offering a promising approach for finding genes with cell-specific expression in less well-studied organisms.

## Methods

### Study design

We isolated cells from five 16-cell stage *Ciona* embryos, each on a different day (Supplementary Table 2). Early ascidian embryos are thought to be bilaterally symmetrical, but to avoid any potential bilateral variation in our small sample, we collected eight cells from the right side of each embryo. The cells were collected individually in batches of eight cells from the same embryo on the same day, with sequencing libraries prepared in parallel, barcoded and then sequenced together. This means that biological variation between embryos and technical variation between batches cannot be distinguished. The advantage of this design is that it minimizes technical variation between cell types of the same embryo and controls for confounding technical and biological variation between embryos. Averaging across the cell types of different batches reduces this unwanted variation, maintaining cell-specific variation. Our results show that cells from the same embryo are more similar to each other than the same cell types are across individuals, with a similar number of genes detected per cell type (Fig. 2a-d).

### Preparation of *Ciona* embryos

*Ciona intestinalis* type A, recently designated *Ciona robusta*^40,41^, adults were obtained from Maizuru Fisheries Research Station (Kyoto University) and Misaki Marine Biological station (The University of Tokyo) under the National Bio-Resource Project for *Ciona*. They were maintained in aquarium in our laboratory at Okinawa Institute of Science and Technology Graduate University under constant light (Calcitrans, Nisshin Marinetech Co., Ltd.) for three days apart from a few hours of darkness a day with feeding to induce spawning of the old eggs. After this, the *Ciona* were maintained under constant light to induce oocyte maturation. Eggs and sperm were obtained surgically from the gonoducts. Embryos were dechorionated after insemination using a solution of 0.07% actinase and 1.3% sodium thioglycolate. Eggs were reared to reach the 16-cell stage in Millipore-filtered seawater (MFSW) at about 18 °C. Embryos from each insemination batch were kept to check the ratio that developed into morphologically normal tailbud. We only used embryos from batches where more than 70% developed normally to tailbud (10 hours post fertilization at 18 degrees) (see Supplementary Table 2 for embryo batch information).

### Naming of cells

In *Ciona*, cells are named using the nomenclature of Conklin^23^: the animal side is prefixed with a lowercase letter (a or b) and the vegetal with an uppercase letter; the anterior with A or a and the posterior with B or b. The initial letter is followed by a number that indicates the embryo stage since fertilization, with individual cells numbered according to their lineage. At the 16-cell stage, the animal domain corresponds to a5.3, a5.4, b5.3 and b5.4, the vegetal domain to A5.1, A5.2, B5.1 and B5.2, and postplasmic RNAs are localized to B5.2.

### Isolation of single cells at the 16-cell stage

At a defined point in development of the 16-cell embryo i.e., at the stage immediately after compaction of the embryo (2.5 ~ 2.6 hours post fertilization), the embryo was transferred to 4°C to slow its development. Each blastomere was isolated with a fine glass needle in a mannitol solution (0.77 M mannitol: MFSW, 9:1) under a stereo microscope at 4 °C regulated by a thermo plate (Tokai Hit Co., Ltd.) and its identity noted. Isolated blastomeres were picked up and transferred immediately with a mouth pipet into a lysis buffer^42^ for reverse transcription.

### Library preparation

We followed the single-cell library preparation method of Tang et al^42,43^ with some modification. We added ERCC spike-in RNA (Thermo Fisher scientific, 4456740, 1:80000) to each lysis buffer and applied 14 and then 9 cycles of PCR amplification after second strand synthesis. Amplified cDNA was purified with MinElute PCR Purification kit (28006, QIAGEN) and QIAquick PCR Purification Kit (28106, QIAGEN) after each PCR reaction respectively and its concentration measured with Qubit^®^ 2.0 Fluorometer (Q32866, Life Technologies) to have more than 150 ng total yield of cDNA. The quality of the amplified cDNA and distribution of DNA fragment size were confirmed by Agilent 2100 Bioanalyzer (Agilent Technologies) with High Sensitivity DNA Kit (5067-4626, Agilent) to consist mainly of 1.0-1.5 kb fragments.

Amplified cDNAs were sheared using sonication Covaris S2 System to produce DNA of 300 bp on average. The settings were as follows: Duty cycle: 20%, Intensity: 5, Cycles per burst: 200, Power mode - Frequency sweeping, Treatment time: 90 seconds, Temperature: 12°C.

NEB Next^®^ ChIP-Seq Library Prep Master Mix Set for Illumina^®^ (E6240, NEB) was applied to sheared cDNA for preparation of the library for the Illumina platform. NEBNext^®^ Multiplex Oligos for Illumina (E7335, E7500, NEB Next Multiplex Oligos for Illumina, NEB) were combined to introduce an index and adaptor to the double-stranded DNA. After extraction of the 300 bp fraction of adaptor-ligated DNA by E-Gel Size Select 2% Agarose (G661002, Invitrogen), DNA was amplified with individual index primers using PCR with 19 cycles.

The amplified DNA fragment composition was purified with Agencourt AMPure XP twice (A63881, Beckman) and again checked by Qubit (> 60 ng of cDNA in total yield) and by Bioanalyzer to ensure that the fragment size was sharply distributed around 300 bp (on average, about 320 bp with a standard deviation of 40). The concentration of fragments with appropriate index adapters was quantified by KAPA Library Quantification Kits (KAPA Library Quantification Kits, Illumina GA/Universal, KK4825, Genetics) to ensure that the final libraries had adapters for both ends and their concentration was at least 20 pM.

### Data generation and quality checking

Libraries were sequenced on Illumina’s (San Diego, CA) MiSeq benchtop sequencer and Illumina HiSeq 2500. Libraries were prepared with different index primers and sequenced on MiSeq using paired 150 nt reads (No. MS-102-2002, MiSeq Reagent Kit v2) with eight multiplexed samples per run with the standard Illumina protocols. The same libraries were sequenced on an Illumina HiSeq 2500 with 150 bp paired end reads (No. PE-402-4001 and FC-402-4001, TruSeq Rapid Cluster - Paired-End and SBS Kits) with 16 multiplexed samples per lane following standard Illumina protocols. Our results from using HiSeq and MiSeq were similar (Fig. 2c-d, cf. Supplementary Figs 3 and 4).

The resulting reads were aligned using Bowtie^44^ version 2.2.6 to the *Ciona* KH genome assembly^45,46^, downloaded from Ghost (http://ghost.zool.kyoto-u.ac.jp/download_kh.html). Reads were mapped using local alignment (--local), with other settings at their default. We did not trim or filter reads, but instead made use of local alignment to find the optimal match. This had the additional benefit that we did not need to split up reads to handle transcripts spanning more than one intron, as is done, for example, in TopHat^47^. Gene counts were calculated from the resulting alignment files using htseq-count^48^ with the non-stranded option and mode “intersection-nonempty” against the KH gene models (version 2013) downloaded from Ghost.

We assessed our samples for mapping quality. We excluded one embryo from subsequent analysis since it had oligo-dT primer sequence in more than 50% of its read pairs; the remaining four embryos had less than 1% of read pairs affected. All remaining samples mapped well to the genome (Supplementary Table 3) and a uniform number of genes were detected (about 60%), although embryo 1 had noticeably fewer detected genes for some of its cells.

### Pattern discovery

Hierarchical clustering to determine candidate patterns was performed with ClusteringComponents in Mathematica 10.4 with the Agglomerate method and Euclidean distance function. This is equivalent to *hclust* in R with the single linkage method. The Transquartile Range (TQR) was calculated as the difference between the first quartile of the ON cluster and the third quartile of the OFF cluster. The quantile method for the TQR used linear interpolation equivalent to type 5 in the R *quantile* function (the hydrologist method). The resulting patterns and TQR score are listed in Supplementary Table 7.

### Single-cell qPCR analysis

cDNA was reverse transcribed from all cells of one embryo per gene replicate using the same protocol we used for single-cell RNA-seq^42,43^. Quantitative PCR was performed using a StepOnePlus PCR machine (Applied Biosystems) with the SYBR green method (No. RR820B, Takara). Each gene was measured with either two or three replicates, except KH.L152.12, which had four. We did not get reliable measurements for KH.S1497.1. The qPCR measures for the cell types of each embryo were scaled between 0 and 1 and then averaged for each cell type across replicates. If there was insufficient target mRNA, it was first amplified using primers covering a wider region of the target gene than those used for single-cell qPCR. Amplification of a specific product in each reaction was confirmed by determining a dissociation curve and comparing with the relevant standard plasmid to estimate the copy number in each cell type. The primers for single-cell qPCR analysis, the IDs for the cDNA clones and the resulting data are listed in Supplementary Tables 4-6.

### In situ hybridisation

Whole-mount in situ hybridisation was carried out as previously described with minor modification^49^. Dig-labeled antisense RNA probes were synthesized in vitro from cDNAs from the *Ciona* cDNA project^50^. mRNA expression was visualized using the NBT/BCIP system (Roche, No.11681451001) and detected on a Zeiss Axio Imager Z1 microscope using DIC (Differential Interference Contrast). The images were acquired with Axiovision SE64 release 4.9.1. Contrast and brightness were adjusted for some images using Adobe Photoshop. The IDs for the cDNA clones are shown in Supplementary Table 5.

### Microarray processing

Previously published microarray data^27^ was processed with the limma R package^51^. Background was corrected using *normexp* and arrays were normalised with the *quantile* method.

### Gene models and names

Gene names for the KH 2013 gene models were downloaded from Ghost (http://ghost.zool.kyoto-u.ac.jp/TF_KH.html and http://ghost.zool.kyoto-u.ac.jp/ST_KH.html) and supplemented with names from Prodon et al^38^.

## Code and Data Availability

RNA-seq data have been deposited in the ArrayExpress database at EMBL-EBI (www.ebi.ac.uk/arrayexpress) under accession number <u>E-MTAB-6117</u>. Software is available at https://github.com/ilsley/Ciona16.

## Acknowledgements

We thank the staff of the Maizuru Fisheries Research Station of Kyoto University and Misaki Marine Biological Station of the University of Tokyo for collecting and cultivating *Ciona* under the National BioResource Project (NBRP) of MEXT, Japan, and RIKEN BRC for providing *Ciona* EST clones through the NBRP. We thank Vladimir Benes and Dinko Pavlinic in the Genomics Core Facility at the European Molecular Biology Laboratory (EMBL) for initial advice on the library preparation protocol and the members of the OIST DNA Sequencing Section for their support in running our samples on their Illumina MiSeq and HiSeq machines. We also thank Sylvain Guillot for his technical support, Filipe Tavares-Cadete for early feedback on the method and Kenji Kobayashi for a helpful comment on the validation experiment. This work was supported by core funding from OIST to the Genomics & Regulatory Systems and Marine Genomics Units.

## Author contributions

NML and NS established and supervised the project. All authors contributed to the study design. RS and TN optimized the experimental protocols, collected and prepared the samples and sequencing libraries. RS designed and performed the in situ and qPCR analysis. GRI conceived and designed the normalization, gene testing and pattern discovery method and performed the bioinformatics analysis. GRI, RS and NML wrote the paper and all authors edited and approved the final manuscript.

## Competing interests

The authors declare no competing interests.

